# Copy parents or follow friends? Juvenile foraging behaviour changes with social environment

**DOI:** 10.1101/429688

**Authors:** Victoria R. Franks, John G. Ewen, Mhairi McCready, Rose Thorogood

**Affiliations:** Department of Zoology, University of Cambridge, Downing Street, Cambridge, CB2 3EJ, U.K.; Institute of Zoology, Zoological Society of London, Regent’s Park, London, NW1 4RY, U.K.; Helsinki Institute of Life Science (HiLIFE), University of Helsinki, Finland; Research program in Organismal and Evolutionary Biology, Faculty of Biological and Environmental Sciences, University of Helsinki, Finland

## Abstract

The first few months of juvenile independence is a critical period for survival as young must learn new behaviours to forage efficiently. Social learning by observing parents (vertical transmission) or others (horizontal/oblique transmission) may be important to overcome naivety, but these tutors are likely to differ in their reliability due to variation in their own experience. How young animals use different social information sources, however, has received little attention. Here we tested if wild juvenile hihi (*Notiomystis cincta*, a New Zealand passerine) retained foraging behaviours learned from parents, or if behaviour changed after independence in response to peers. We first trained parents with feeders during chick rearing: one-third could access food from any direction, one-third could access food from one side only, and the remaining third had no feeder. During post-fledge parental care, juveniles chose the same side as their parents. Once independent, juveniles formed mixed-treatment groups naturally so we then presented feeders with two equally profitable sides. Juveniles with natal feeder experience were quicker to use these feeders initially, but side choice was now random. Over time, however, juveniles converged on using one side of the feeder (which differed between groups). This apparent conformity was because juvenile hihi paid attention to the behaviour of their group and were more likely to choose the locally-favoured side as the number of visits to that side increased. They did not copy the choice of specific individuals, even when they were more social or more familiar with the preceding bird. Our study shows that early social experiences with parents affect foraging decisions, but later social environments lead juveniles to modify their behaviour.

## Introduction

The first few months of independent life are a critical period for survival in many bird species. Studies have reported that only 50% of young survive the first two months after leaving parents (Cox *et al*., 2014; Naef-Daenzer and Grüebler, 2016), and in some populations first-year survival can be as low as 11% (Sullivan, 1989; 30% first-year survival reported by McKim-Louder *et al*., 2013). Mortality rates often peak over winter and can reach five times that of adults as juveniles struggle to survive in harsh environmental conditions (Goss-Custard and Durell, 1987; Daunt *et al*., 2007). What determines individual survival is likely to be non-random (Naef-Daenzer and Grüebler, 2016), and starvation plays a key role (Ringsby, Sæther and Solberg, 1998; Sol *et al*., 1998; Daunt *et al*., 2007; Low and Pärt, 2009). However, little is known about how young birds learn to find food and survive during their first few months (Cox *et al*., 2014).

One general problem for young animals is their inexperience compared to adults (Galef and Laland, 2005). This means they have had limited opportunities to learn to find, capture, and process food (Marchetti and Price, 1989; Wheelwright and Templeton, 2003), which can limit their foraging efficiency. For example, juvenile garter snakes (*Thamnophis atratus hydrophilus*) feed on a more restricted range of food types than adults (Lind and Welsh, 1994), and studies in young birds show they take longer to forage than adults (Daunt et al., 2007; Gochfeld and Burger, 1984; Kendal et al., 2009; Marchetti and Price, 1989; Sol et al., 1998). Therefore, juveniles need to learn new behaviours to improve their efficiency. However, learning also presents a challenge to juveniles and they may take longer than adults to acquire new skills (Franks and Thorogood, 2018). If young animals face both a greater need to learn and an increased cost of learning, are there strategies they can use to overcome these combined challenges? Paying attention to the behaviour of others can be one way for juveniles to buffer their own inexperience (Galef and Laland, 2005; Kendal *et al*., 2005; Kitowski, 2009; Clutton-Brock, 2016; Griesser *et al*., 2017) and young animals encounter a variety of sources of social information during their first few months.

Before they become fully independent, naïve juveniles can learn important behaviours from their parents (“vertical transmission” (van Schaik, 2010)), such as preference for or aversion to certain foods (Galef and Giraldeau, 2001), or foraging techniques (Rapaport, 2006; Geipel *et al*., 2013). In some cases, experiences with parents have long-term effects on behaviour later in life. For example, cross-fostered blue tits (*Cyanistes caeruleus*) and great tits (*Parus major*) shifted their foraging niche in the direction of their foster parents and maintained this preference to adulthood when feeding their own young (Slagsvold and Wiebe, 2011). Reliable vertical social learning is in the best interests of both parents and juveniles, because it increases offspring survival and maximises lifetime reproductive fitness (Clutton-Brock, 1991; Laland and Kendal, 2003; Thornton and Clutton-Brock, 2011). However, if parental information is less optimal (Farine, Spencer and Boogert, 2015) or environments change so that behaviours learned in early life become outdated (Wong and Candolin, 2014), then young animals should pay attention to other information to update behaviour.

Once independent, juveniles encounter other individuals (“peers”). When peers are present in the same location, at the same time, and encounter the same environment as the naïve individual, learning via other juveniles (horizontal transmission”) and adults (“oblique transmission”) (van Schaik, 2010) may provide more up-to-date information. Under some conditions, animals rely on copying the behaviour of peers to such an extent that one behaviour becomes predominant in a group, leading to conformity across individuals (van de Waal, Borgeaud and Whiten, 2013; Aplin *et al*., 2015b). However, peer-provided information could conflict with previous information from parents and can also be unreliable for a variety of reasons. Peers may provide deliberately misleading information: for example, fork-tailed drongo (*Dicrurus adsimilis*) use false alarm calls to scare others away from food (Flower, 2011; Flower, Gribble and Ridley, 2014). Peers can also learn incorrectly, so juveniles risk copying maladaptive behaviours (Curio, Ernst and Vieth, 1978; Laland and Williams, 1998; Franz and Matthews, 2010). Finally, animals have preferred and avoided companions, forming “social networks” which determines what peers and social learning opportunities they encounter (Krause, Lusseau and James, 2009; Kurvers *et al*., 2014; Krause *et al*., 2015). Variation in familiarity can influence social information use, although sometimes animals may prefer to learn from familiar partners (Swaney *et al*., 2001; Guillette, Scott and Healy, 2016), while others use unfamiliar sources which have different personal experiences (Ramakers *et al*., 2016). Further, individuals that interact with many peers (have a high “degree”) encounter many sources of information, and may acquire behaviour faster or gain a more complete picture of the environment (Aplin *et al*., 2012; Tóth *et al*., 2017).

Both theory and empirical studies have discussed how animals adjust their use of social and personal information to best suit current conditions (Kendal, Coolen and Laland, 2004; Kendal *et al*., 2005; Thorogood and Davies, 2016), but less is known about how they trade off or integrate different sources of social information from parents and peers (Laland, 2004; Farine, Spencer and Boogert, 2015). This may be important for understanding how behaviours learned early in life persist, particularly in social groups where animals do not associate randomly. Hihi (*Notiomystis cincta*), a threatened New Zealand passerine, provide an ideal opportunity to investigate social information use in young wild birds. In one population (Tiritiri Matangi Island), hihi nest and raise their altricial chicks in monitored nest-boxes in territories during the breeding season (September-February). Fledglings are cared for by parents for two weeks before dispersing from the nest site, and then form groups of independent juveniles in reliable locations on the island. The time with parents and in groups likely provides opportunities for social learning, but how these different sources of information are used has not been investigated. Finally, understanding the importance of social learning for foraging in young hihi may help us understand how they adjust feeding behaviour following conservation interventions, particularly when provisioning supplementary food is a crucial part of conservation management for hihi (Cox *et al*., 2014).

To test the hypothesis that social experiences in early life affect foraging behaviour of juvenile hihi, we set up novel feeders at nests and at sites where groups congregate. We predicted (1) young hihi use social information provided by parents during their first couple of weeks post-fledging; and (2) this information continues to influence their behaviour once independent. However, if (3) juveniles pay more attention to social information in groups, then their behaviour would change once independent and depend on social characteristics (tie strengths and degree). Finally, to highlight how the inexperience of juveniles changes their learning strategies we also predicted (4) juvenile hihi respond to social information more than adults. By recording sequential visits to feeders, we could detect copying and changes in behaviour through time.

## Methods

### Study Population

We conducted our experiment during one breeding season (October 2015 – April 2016) on Tiritiri Matangi Island (36°36’00.7”S, 174°53’21.7”E). The study population of hihi numbered c. 88 adults and 132 juveniles (juveniles: any fledgling from the 2015–2016 breeding season, adults: all other birds) (McCready and Ewen, 2016). Each individual was identifiable from a unique combination of coloured leg rings. During our study hihi also carried a Radio Frequency Identification (RFID) Passive Integrated Transponder (PIT) tag (from here, “PIT tags”) integrated into one of the leg rings (IB Technology). This enabled remote recording of visits to feeding stations fitted with antenna and data-loggers (IB Technology model EM4102).

### (1)Learning with parents at nests

#### Experimental procedure

We divided active nests into three treatment groups (Figure 1a): “no feeder” (naïve); an “open feeder” which could be entered from any direction so hihi learned to recognise feeders as a food source; and a “side-choice feeder”, where only one of two channels (left, LHS or right, RHS, assigned equally among nests) contained a sugar water reward so hihi learned an association and a side choice. These feeder treatments ensured that fledglings had different experiences of feeders with parents. Nests were allocated based on their location and surrounding forest maturity, which balanced rearing conditions of chicks and avoided movement between different treatments. Each treatment contained similar numbers of nests and fledglings (Figure 1a). We only included first clutch nests in treatments to avoid fledglings encountering different feeders at any later second-clutch nests once they dispersed from the nest site.

**Figure 1.**
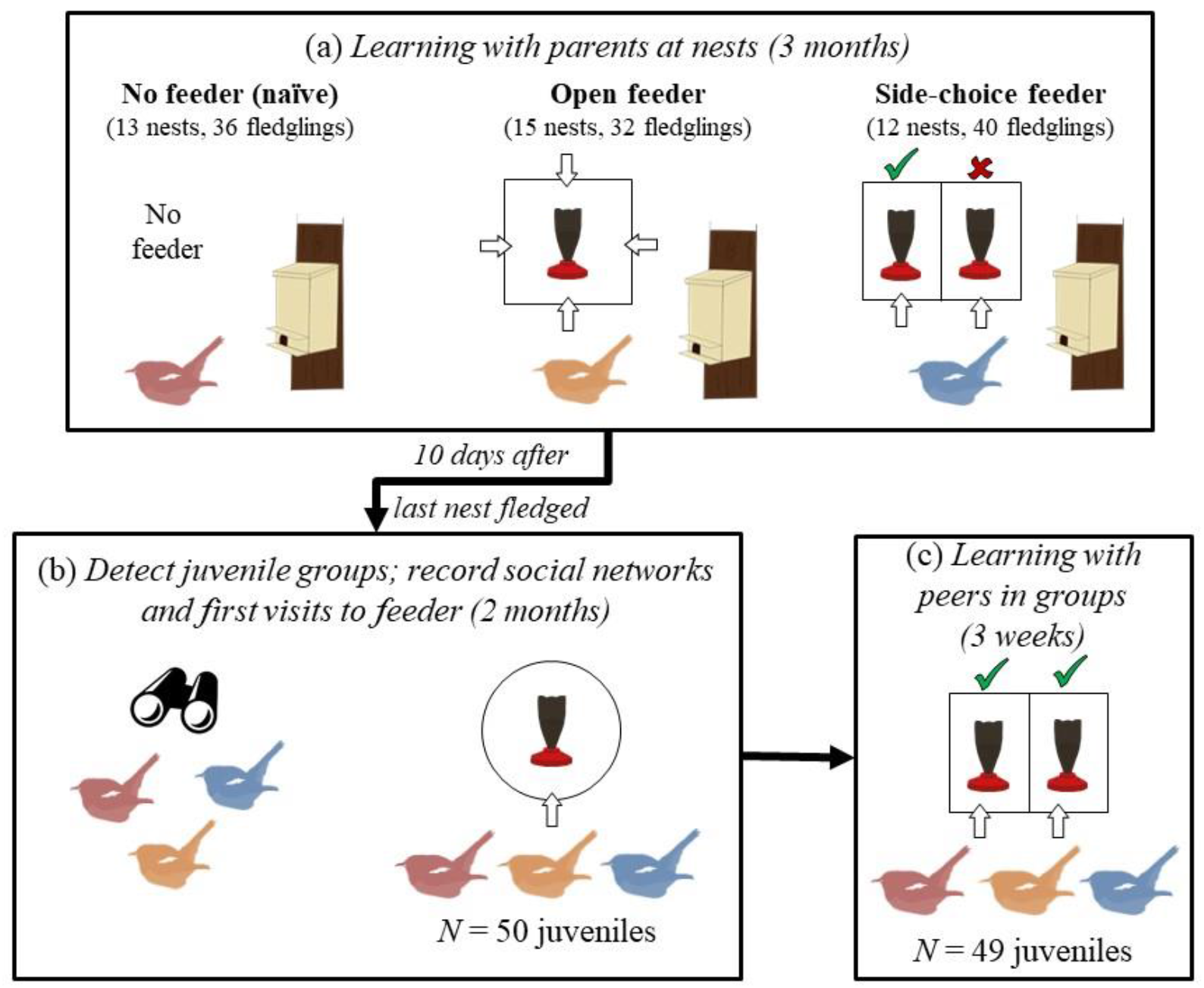
The stages of the experiment: (a) learning with parents at nests, showing feeder designs for each treatment, number of nests and fledglings assigned to each treatment group (no feeder: red, open feeder: orange, side-choice feeder: blue); (b) detecting juvenile group sites (containing a mixture of juveniles from each nest treatment group), then recording visits to a supplementary feeder to assess how quickly juveniles from different treatment groups visited feeders and construct a social network; (c) learning with peers at side-choice feeders once independent and in group sites. Not all juveniles were consistently present throughout all of (b) and (c).

Between November 2015 and January 2016, we set up feeders 10 days after chicks hatched, approximately 10 metres from nest-boxes. This gave parents two weeks to learn to use the feeders before chicks fledged (hihi fledge 28 days after hatching). Sugar water was provided in semi-opaque brown bottles attached to Perky Pet^®^ feeder bases (213 Pop Bottle Hummingbird Feeder) which did not provide hihi with visual information about contents prior to foraging. Feeders were checked daily; if we did not observe at least one parent using the feeder in the first three days, we moved it following protocols used in previous studies (Ewen *et al*., 2008; Thorogood, Ewen and Kilner, 2011). At all side-choice feeders, parents first showed no preference for a side (exactly half of all first visits were to the rewarding side) but then learned to use the reward side before chicks fledged (see supplementary information on parent learning at nests). Parents did not use open feeders at 6 of the 15 nests, but as their fledglings (*N* = 13) still had an opportunity to observe the feeder in the parents’ territory, we retained them in the treatment group for later analyses.

After chicks fledged, we observed visits to all feeders (for 45 minutes each, at least every second day) and also recorded visits to side-choice feeders with RFID data-loggers at entry points. Additionally, we monitored feeder visits to side-choice feeders using Bushnell NatureView HD^®^ trail cameras placed approximately 50cm from the feeder. We accounted for differences between continuous recording at side-choice feeders and the shorter observations at open feeders in later analyses by only considering if we ever recorded use by each fledgling, rather than the frequency or time of use. After each observation period, we located and identified any fledglings additionally heard within a c.15-metre radius of the feeder. We removed nest feeders once fledglings were not seen or heard at the nest site for two consecutive days (suggesting they had dispersed, or died); this occurred on average 10 days after fledging (range 0 – 13 days). At any nests where we never heard or saw fledglings, we waited 10 days before taking down feeders.

#### Data analysis

All analyses (of both parent and peer effects) were conducted in R (version 3.5.0) (R Core Team, 2017). We used a binomial sign test to determine if fledglings made the same choice as their parents when first visiting side choice feeders. We compared the number of instances when fledglings from side-choice feeder nests chose the same side (LHS or RHS) as their parents to an expected random chance of 0.5.

### (2) Learning with peers in groups

#### Detecting juvenile groups, and recording a social network

We started surveying for juvenile groups 10 days after the last nest fledged. Every day for three weeks we recorded ring combinations of all hihi sighted during one-hour surveys conducted in six forested gullies across the island (Figure 1b). We selected two sites c. 300 metres apart (“Site 1” and “Site 2”) where we consistently recorded the most juveniles (results not presented here). These sites were located in latitudinally-orientated valley gullies containing mature remnant forest that were separated by a parallel ridge with open pasture. To record a social network (Figure 1b) before testing for retained responses from nests or for horizontal transmission from peers (Figure 1c), we set up a feeder with one entry point (i.e. no side choice) at each of these two group sites for six weeks. We recorded time-stamped visits with a RFID data-logger at the entry point and collected a total of 11928 visits by 50 juveniles (plus 14 adults that also visited). We used these visits to construct one weighted social network with the function “gmmevents*”* in the R package asnipe (Farine, 2013), which calculates associations based on similarities in timings of visits. Any juveniles that visited fewer than three times (*N* = 3) were excluded. Using this network, we calculated each individual’s degree centrality (number of and strength of associations), and tie strengths between hihi (number of times each pair of hihi in the network associated). Most juveniles present in later stages of the experiment used these “network feeders” (42/50), so by the end of network data collection the majority of juveniles had experienced entering a feeder, but not all had experience of a side-choice design (20/50 from side-choice feeder nests). The 8 juveniles that arrived after network recording did not have degree or tie measures during later analysis of effects of sociality on foraging behaviour. No new hihi were recorded after six weeks, suggesting that the majority of hihi using group sites were included in the network.

#### Learning from peers

Following network recording, we replaced network feeders with our side-choice feeder design; however, now both sides were equally rewarding and contained sugar water (Figure 1c; Figure 2). These feeders tested for retained side preference from nests, or effects of peers on foraging behaviour (side choice). During set-up, we ensured that the location and density of vegetation surrounding the feeder was as similar as possible to limit external influences on side choice. We recorded visits to both sides using an RFID data-logger at each entry point and also placed trail cameras 1 metre from feeders to capture images of visits. We continued observations on alternate days in a 10-metre radius of the feeder (total of 25 one-hour observations per site) and recorded identities of hihi present in 30-second time blocks (120 blocks per survey). These observations were used to indicate how long hihi spent near the feeder when they could be observing others (but not necessarily visiting the feeders and detected by the RFID system). Feeders were set up for three weeks (at Site 2, no visits were recorded for days 14 and 15 because of problems with the RFID data-loggers).

**Figure 2.**
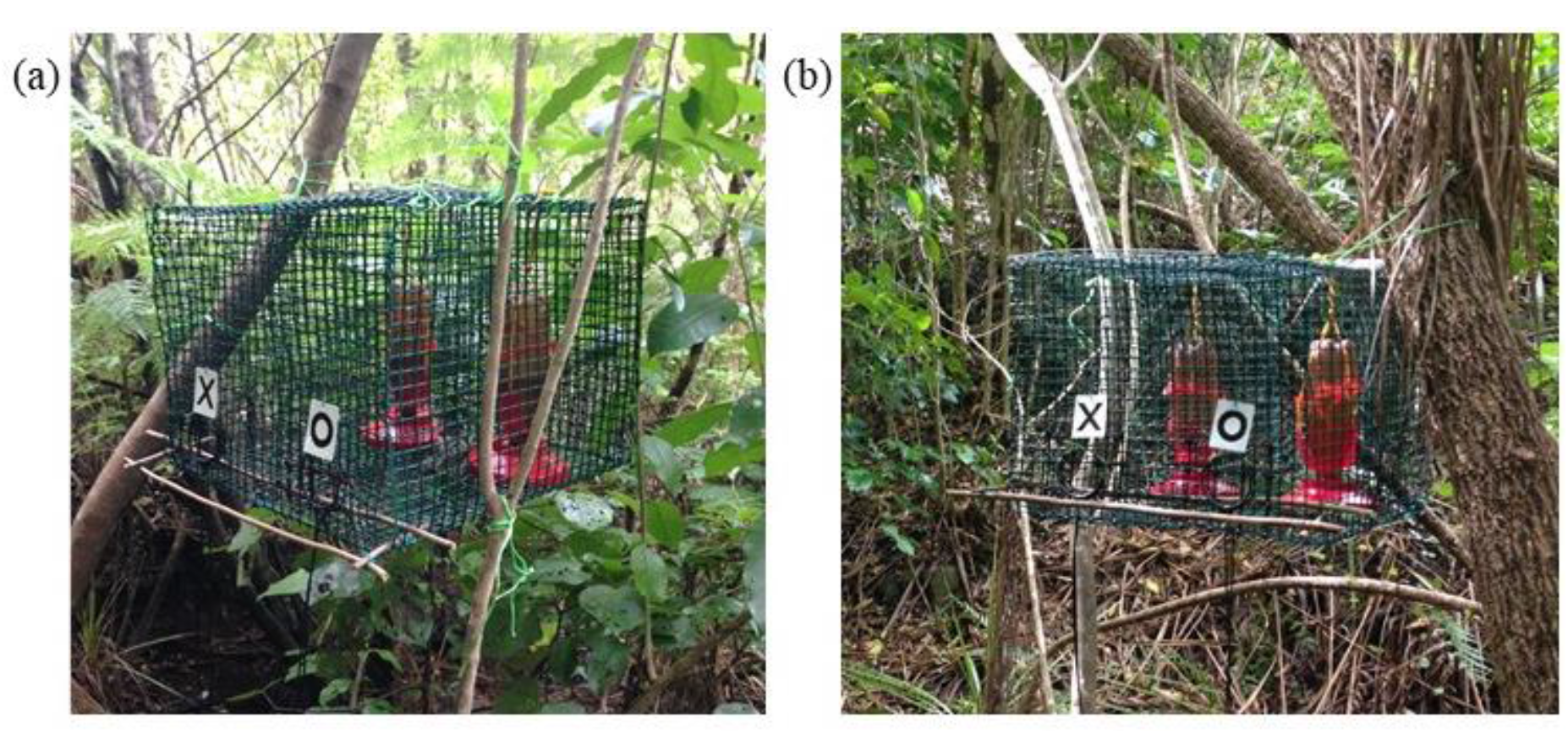
Photos of feeder set-up for juvenile groups in (a) Site 1; (b) Site 2. “X” and “O” symbols were used for another experiment, but “X” is always left, “O” right. Photo credits: Rose Thorogood.

In total across (during surveys, network recording, and feeder presentations), we recorded 62 first-clutch juveniles (no feeder *N* = 15; open feeder *N* = 22; side-choice feeder *N* = 25), although not all individuals were observed during each stage of the experiment. There was no difference in the proportion of fledglings recorded from nests with or without feeders (Fisher’s exact test of fledglings detected at group sites; from nests with feeders = 47/74; from nests without feeders = 15/29; *P =* 0.37). We also recorded 9 second-clutch juveniles. We added these individuals to the naïve treatment in later analyses as there was no evidence they appeared at group sites any later than naïve first-clutch juveniles (compared ranked order of arrival; Wilcoxon rank sum test: *W* = 30, *P* = 0.12).

#### Data analysis

For analyses using Generalised Linear Models (GLMs) or Generalised Linear Mixed Effect Models (GLMMs), we used a model selection approach (Burnham and Anderson, 2002; Symonds and Moussalli, 2011; Harrison *et al*., 2017) in the R package AICcmodavg (Mazerolle, 2017). We ranked candidate Generalised Linear Models (GLMs) or Generalised Linear Mixed Effect Models (GLMMs) that included all possible combinations of relevant predictors by their corrected Akaike Information Criterion (AICc). For all models within 2 AICc units of the top-ranked model, we calculated averaged effect sizes (±95% confidence intervals) of included predictors to assess their effect (Burnham and Anderson, 2002; Nakagawa and Cuthill, 2007). Any effect where confidence intervals did not span 0.00 were considered significant. GLMMs were implemented using the lme4 package (Bates *et al*., 2015).

#### First visits: did juveniles retain behaviour from experiences with parents?

To detect any effects of nest experiences once juveniles were independent, we ranked the times of first visits (latency in seconds from the very first visit, specified as 0 seconds) for all juveniles that visited during feeder presentations (*N* = 58: naïve = 21; open feeder = 16; side-choice feeder = 21), which accounted for non-normally distributed times. This included eight juveniles that only visited after the network feeder (naïve = 5, open feeder = 2, side-choice feeder = 1). We used Poisson-distributed GLMMs to analyse variation in arrival rank depending on whether a feeder was present or absent at nests, and for juveniles from nests provided with feeders, whether they or their parents had used feeders and the feeder type. We included a random intercept to account for whether juveniles arrived for the first time during network recording or experiment feeders.

We tested for a bias in side choice across all juvenile first visits using a Fisher’s exact test to compare the number of LHS and RHS choices by all juveniles to a random distribution within and across sites. We also used a binomial sign test to test whether the subset of juveniles from side-choice nests retained a side preference by comparing the number of times individuals chose their nest reward side to a random choice (50% of visits). Finally, we assessed if first choice was socially-mediated using a binomial GLM to analyse if juveniles chose the same side as the preceding bird (yes = 1, no = 0) depending on how closely they followed that bird in time (log10-transformed seconds between visits because times were not normally distributed).

#### Ongoing visits: did juveniles copy peers?

We investigated changes in side choice by groups over the course of the experiment (mean ± S.E. days individuals were recorded visiting feeders = 11.47 ± 0.80). We calculated the proportions of all visits made per day to the RHS at each feeder (including all visits irrespective of age) and used binomial GLMs to test if changes in daily group preference depended on experiment day and group site. Following the results from this initial analysis, we began investigating individual level patterns. To test if juveniles from side-choice nests continued to prefer their nest reward side, we calculated the proportions of visits each side-choice nest juvenile made per day to the RHS. We used GLMMs to investigate if this proportion changed across experiment days and depending on nest side (RHS or LHS) between the two group sites. Preference would differ between juveniles across days if side choice resulted from nest treatment, so we included an interaction between nest side and experiment day. Our random intercept was individual identity to account for repeated data-points for the same birds. To further explore that juveniles from opposite side-choice feeder nests were mixed between the two groups and did not drive changes in side preferences, we compared the proportion of each juvenile’s total visits made in Site 1 between RHS and LHS side-choice nest juveniles, using a Wilcoxon rank sum test.

Juvenile groups contained a mix of birds from different nest treatments (Figure 1), so we analysed if hihi copied (i) the behaviour of the social group and (ii) the behaviour of specific individuals, on each visit. For (i), we analysed if hihi chose the side favoured by each group by the end of the experiment (the “locally-preferred side”: Site 1 = RHS, Site 2 = LHS), depending on group preference. For each visit, we calculated the group’s preference as the frequency of preceding visits made that day to the locally-preferred side. Although frequency of behaviour may not represent the preference of all individuals if some individuals visit more often (Aplin *et al*., 2015a; van Leeuwen *et al*., 2015, 2016; Whiten and van de Waal, 2016), we initially calculated frequency of behaviour and frequency of individuals and they were strongly correlated (Pearson correlation: *r* = 0.81, *P* < 0.001). Thus, we used frequency of behaviour as our measure of group preference, which does not require hihi to recognise and track individuals (Aplin *et al*., 2015a, 2015b). Binomial GLMMs were then used to test if an individual’s side choice at each visit matched the locally-preferred side (1), or did not match (0) depending on group preference, or if there were effects of day of experiment (days 1–21, to assess if side choice varied more with social environment or personal learning (Aplin *et al*., 2015a, 2015b)), time of day (hours; individuals visiting later in the day could have observed more visits), the focal bird’s degree score from the network (as a measure of the effects of sociality on behaviour), and age (to test if juveniles used social information differently to adults).

To explore if (ii) hihi copied specific individuals, we calculated the proportion of times each individuals’ side choices were matched by the bird that visited next. We then used binomial GLMMs to explore if individuals that visited closer in time to the preceding bird (log10-transformed seconds to account for non-normally distributed times) were more likely to choose the same side (no = 0, yes = 1). Including time allowed us to explore if temporal proximity allowed for stronger copying, or conversely if closely-following individuals avoided each other to limit competition for resources (Krebs and Inman, 1992). We also included additional fixed effects of tie strength (familiarity) and focal individual degree (sociality) to investigate social effects on copying specific individuals, and age of both the preceding hihi and the focal individual (age could affect social information use). Finally, we included measures of group preference per visit, to assess how copying specific individuals affected side choice in addition to any effects from the broader social environment. As a random intercept, we included individual identity to account for repeated visits by individuals.

Most juveniles moved between group sites at least once (32/49 juveniles, but only two adults), so we repeated analyses for (i) and (ii) to test if limited or outdated personal information affected copying after changing sites in juveniles (Laland, 2004; Kendal *et al*., 2005). Explanatory variables were the same as in the first sets of analyses (excluding age), but we also included number of site changes (because earlier experiences could affect side choice). For analysis of choice of locally-preferred side depending on the behaviour of the group, we additionally included the proportion of visits each juvenile made to the preferred side at the previous site on the previous day, as individuals with a stronger preference for a side may have been less likely to switch sides after moving.

## Results

### Learning with parents at nests

Only fledglings of parents that used feeders themselves (at 60% of open and 100% of side-choice feeders) were detected using feeders at nest sites (open: 7/32 fledglings at 4/15 nests; side-choice: 11/40 fledglings at 5/12 nests). At nests where parents did not use feeders some fledglings were not observed around the nest site after fledging (N = 6 nests, 13 fledglings), but this was not only due to mortality as 9/13 were recorded once they were independent. Where we did not observe any feeder use, we were confident hihi did not use feeders at other times as there was no sugar water taken or residue left behind on feeder bases (which we saw at feeders with confirmed use).

The majority of the 11 fledglings that visited side-choice nest feeders (5 nests) chose the same side as their parents on their first visit (9/11 used same side; Binomial sign test: *P* = 0.03). At all five nests, we observed fledglings follow a parent into the vicinity of the feeder while begging (Figure 3a, b), and then follow a parent into the feeder (often still begging) (Figure 3c). Five fledglings visited feeders only once; in the remaining six, their number of visits ranged from 3 – 51. Five out of six fledglings maintained a preference for their side but occasionally also visited the other side (mean = 74% preference).

**Figure 3.**
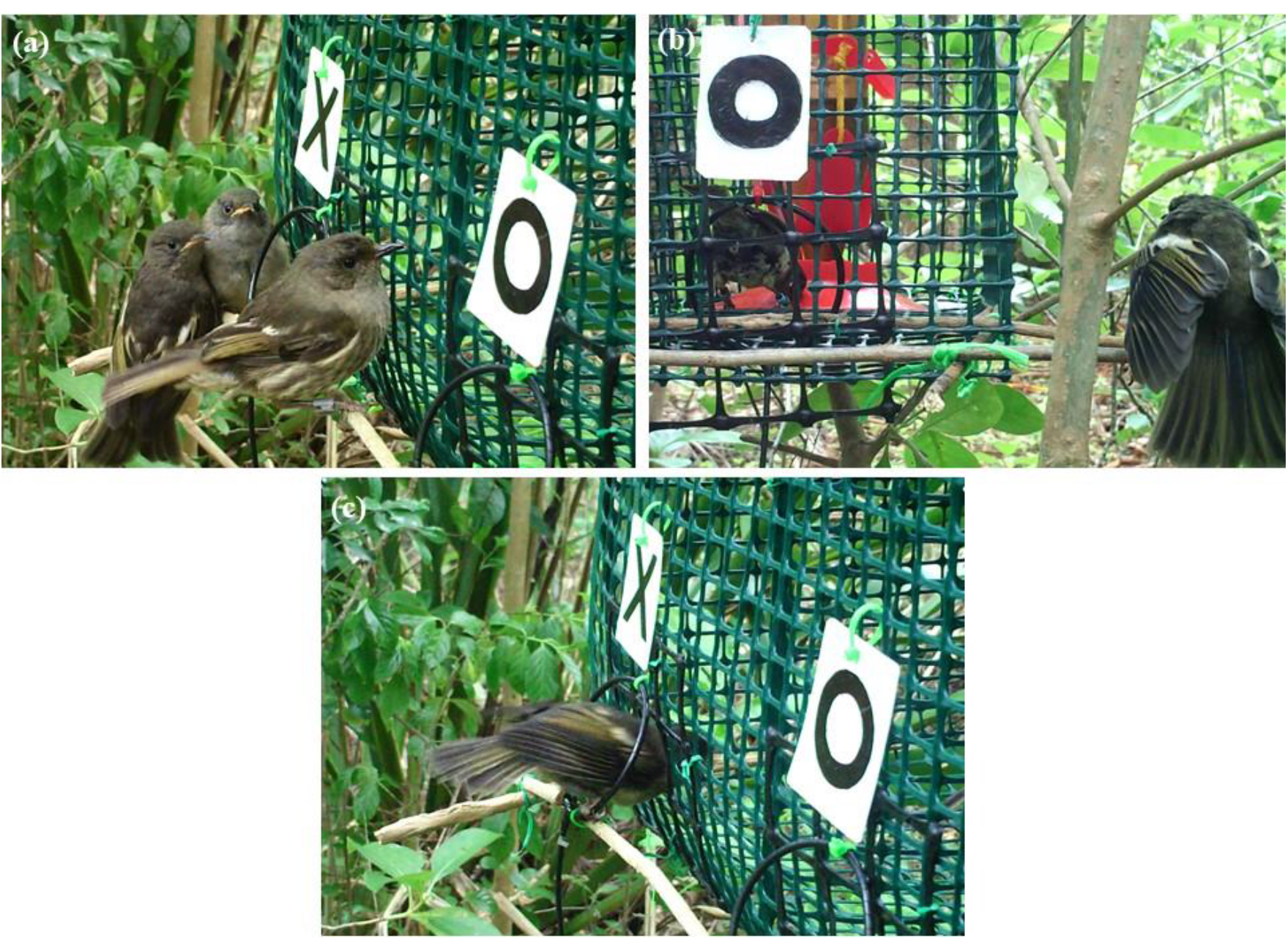
Behaviour of fledglings with parents at side-choice feeders. (a) Two fledglings (left) follow mother (right) to feeder; (b) a fledgling (right) in begging posture, while parent is in feeder; (c) fledgling enters feeder following a parent (out of frame on right). “X” and “O” symbols were used for another experiment, but “X” was always left, “O” right. Images captured using trail cameras.

### Learning with peers in groups

#### First visits: did juveniles retain behaviour from experiences with parents?

Juveniles from nests with feeders were quicker to use group site feeders than juveniles from nests with no feeder (*N* = 58; 9/10 of the first juveniles were from nests with feeders; effect of feeder presence on arrival rank = –0.27±0.05, 95% CI = –0.37 – –0.18; Figure 4; Supplementary Table 1a). However, the details of early life experiences were not important for juveniles from nests with open and side-choice feeders: models containing parents’ use of the feeder, the type of feeder, and if a juvenile had used its nest-site feeder were ranked lower than the null model (Figure 4; Supplementary Table 1b). Furthermore, juveniles from side-choice feeders did not significantly prefer the side experienced at the nest (binomial sign test, 10/16 juveniles chose nest side on first visit, *P* = 0.46), even when they had used nest feeders themselves (binomial sign test, 5/8 chose nest option on first visit, *P* = 0.73).

**Figure 4.**
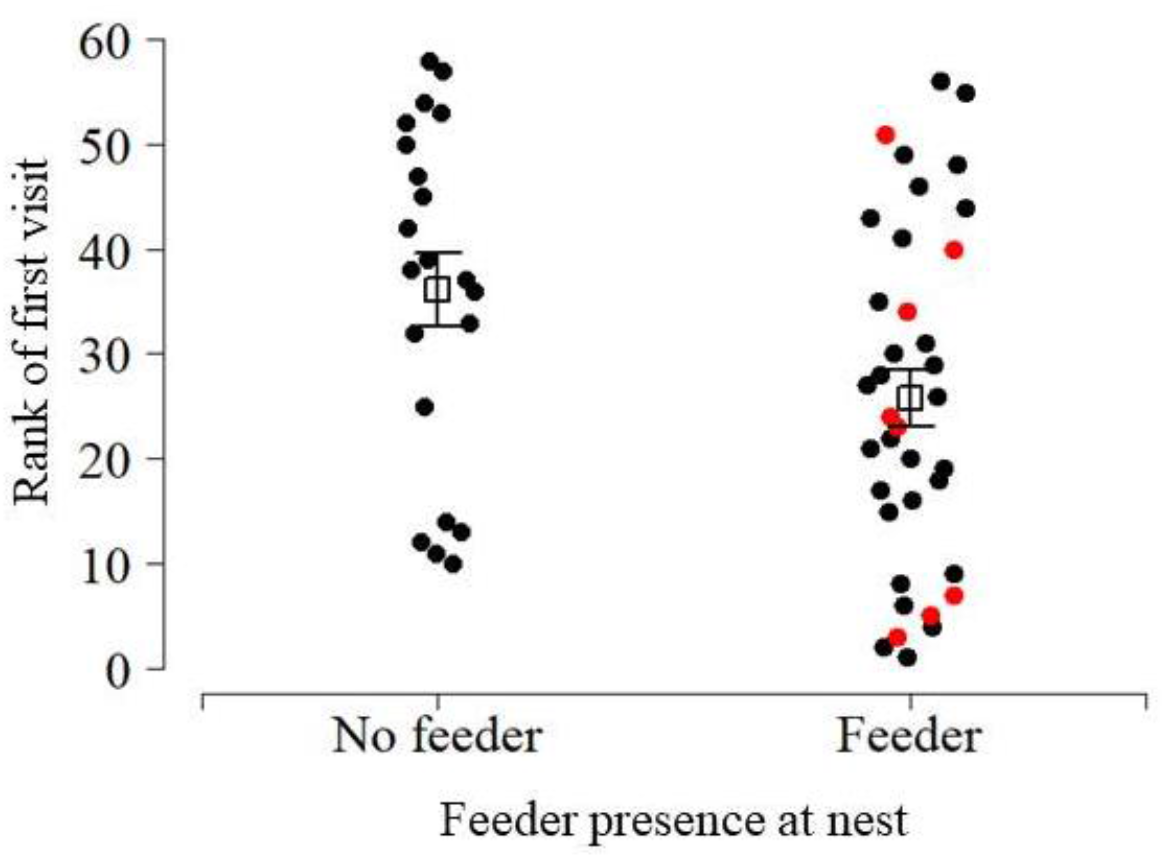
Ranks of first visits to group site feeders for juveniles from nests with no feeder and nests with feeders. Red points depict juveniles from nests where parents did not use feeders. Open squares depict mean rank (± standard error).

Across their first visits juveniles showed no preference for a side within sites, and preference also did not differ between sites (Fisher’s exact test: Site 1 = 10/24 visits to RHS, Site 2 = 15/25 visits to RHS, *P* = 0.26). For comparison, adults also showed no preference at either site (Fisher’s exact test: Site 1 = 2/12 visits to RHS, Site 2 = 6/12 visits to RHS, *P* = 0.19; 4 parents [3 males, 1 female] from side-choice feeder nests visited, but none chose their nest side on first visit). Peers also had little influence on side choice during first visits. Most juveniles first visited on day 1 or 2 (35/49, 71%), and half were within 2 minutes of the previous hihi (22/47 47%; the first visitors to each site are excluded). However, hihi only chose the same side as the previous bird 45% of the time (21/47) and were not more likely to copy if visits were closer together (null model ranked higher than one including latency from previous bird’s visit; Supplementary Table 2).

#### Ongoing visits: did juveniles copy peers?

Despite both sides being equally rewarding, hihi groups developed a local preference for one side as the experiment progressed, but in the opposite direction at the two sites (*N* visits = 10049; the only model with ΔAICc < 2 included experiment day*site; effect = –0.08±0.01, 95% CI = –0.09 – –0.07; Figure 5a, b; Supplementary Table 3a). This was not because of assortment of juveniles from the different side-choice nests as they did not differ in their proportion of visits at the two sites (no difference in proportion of visits juveniles from each group made at Site 1; Wilcoxon rank sum test: *W* = 33.5, *P* = 0.87) and did not maintain a significant preference across their visits within each site (null model ranked highest; Supplementary Table 3b). In all birds, side chosen on first visit did not predict a hihi’s overall side preference in either site (Fisher’s exact test: Site 1: 23/36 hihi maintained a preference for the same side chosen on first visit; Site 2: 23/38 maintained a preference; *P* = 0.81). For all juveniles (*N* = 49), but not adults (*N* = 25), side choice was best explained by the strength of the group’s preference that day for the locally-preferred side (age*social environment, Table 3a; Figure 5c; Supplementary Table 4a). By comparison, adults developed a stronger preference over days compared to juveniles (visit day*age, Table 3; Supplementary Table 4a). There was a non-significant trend for side choice in both adults and juveniles to follow the group’s preference more strongly later in the day (Table 3a; Supplementary Table 4a), but there was no evidence that having more associates affected side choice (no effect of degree on choosing locally preferred side: Table 3a; Supplementary Table 4a).

**Figure 5.**
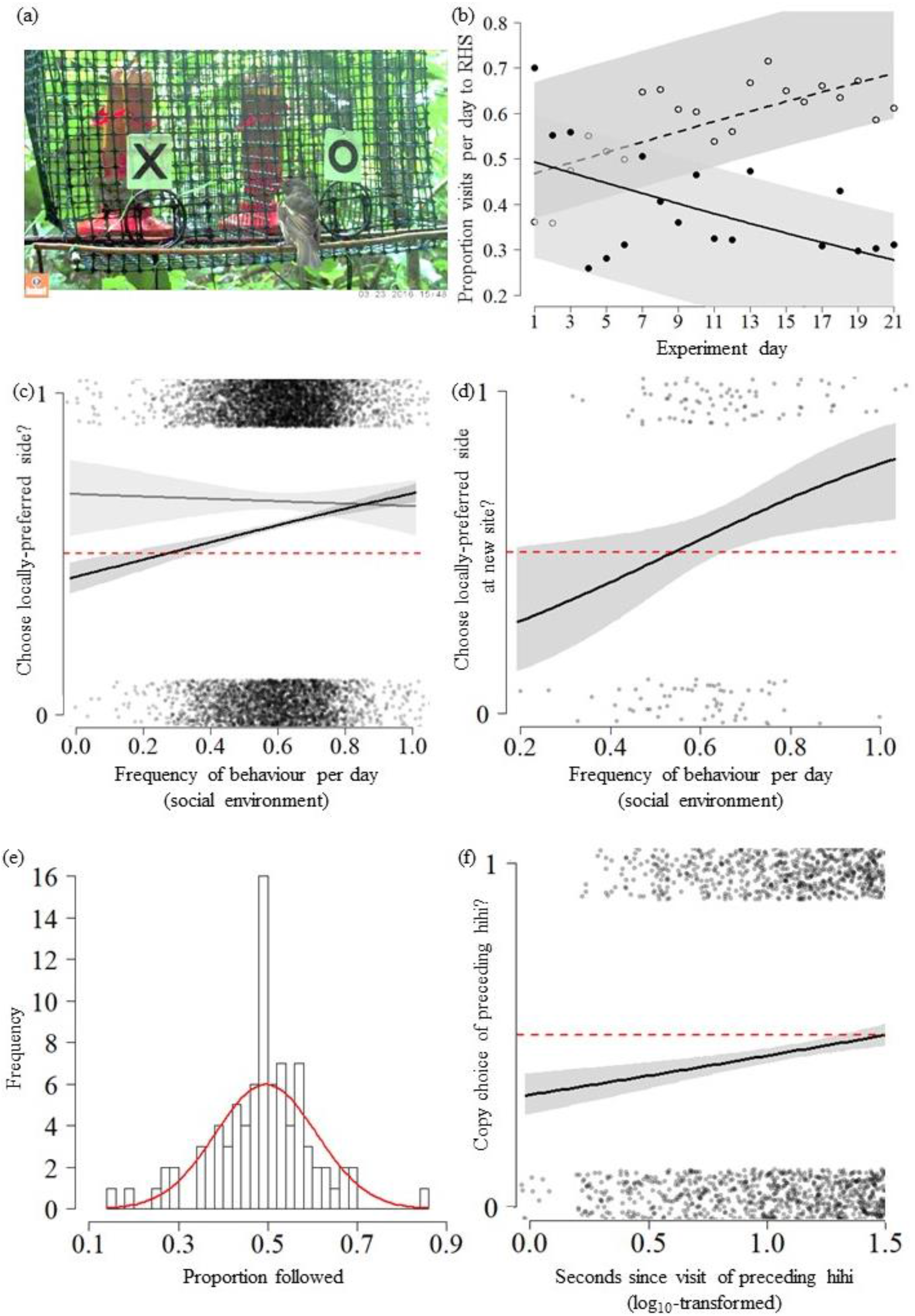
(a) juvenile hihi visiting group site feeder; (b) proportion of visits recorded per day to RHS at group feeders (Site 1 = dashed line; Site 2 = solid line), which shows how preference for one side of the feeders changed across the experiment; (c) likelihood that juveniles (black line) and adults (grey line) chose the local side (yes = 1, no = 0), depending on frequency its use by other hihi that day. Shows that side choice depended on social group preference in juveniles, but not adults; (d) likelihood that juvenile hihi (black line) chose the local side when they changed between sites (yes = 1, no = 0), depending on frequency its use by other hihi that day. Shows that juveniles copied group preference when they moved to a new site; (e) Frequencies of the proportion of visits by each individual where their side choice was copied by the following hihi, where red line represents normal distribution for reference. Shows that most individuals were only copied by the next bird on 50% of visits; (f) likelihood that hihi chose the same side as the previous bird (yes = 1, no = 0), for visits under 30 seconds apart. Shows that hihi were less likely to copy the preceding bird when they visited very soon after. All predicted model estimates and 95% confidence intervals (grey areas) come from top-ranked models. Points are jittered to improve visibility, so do not show exact values. Dashed red lines indicate 50% likelihood of copying.

**Table 3.**
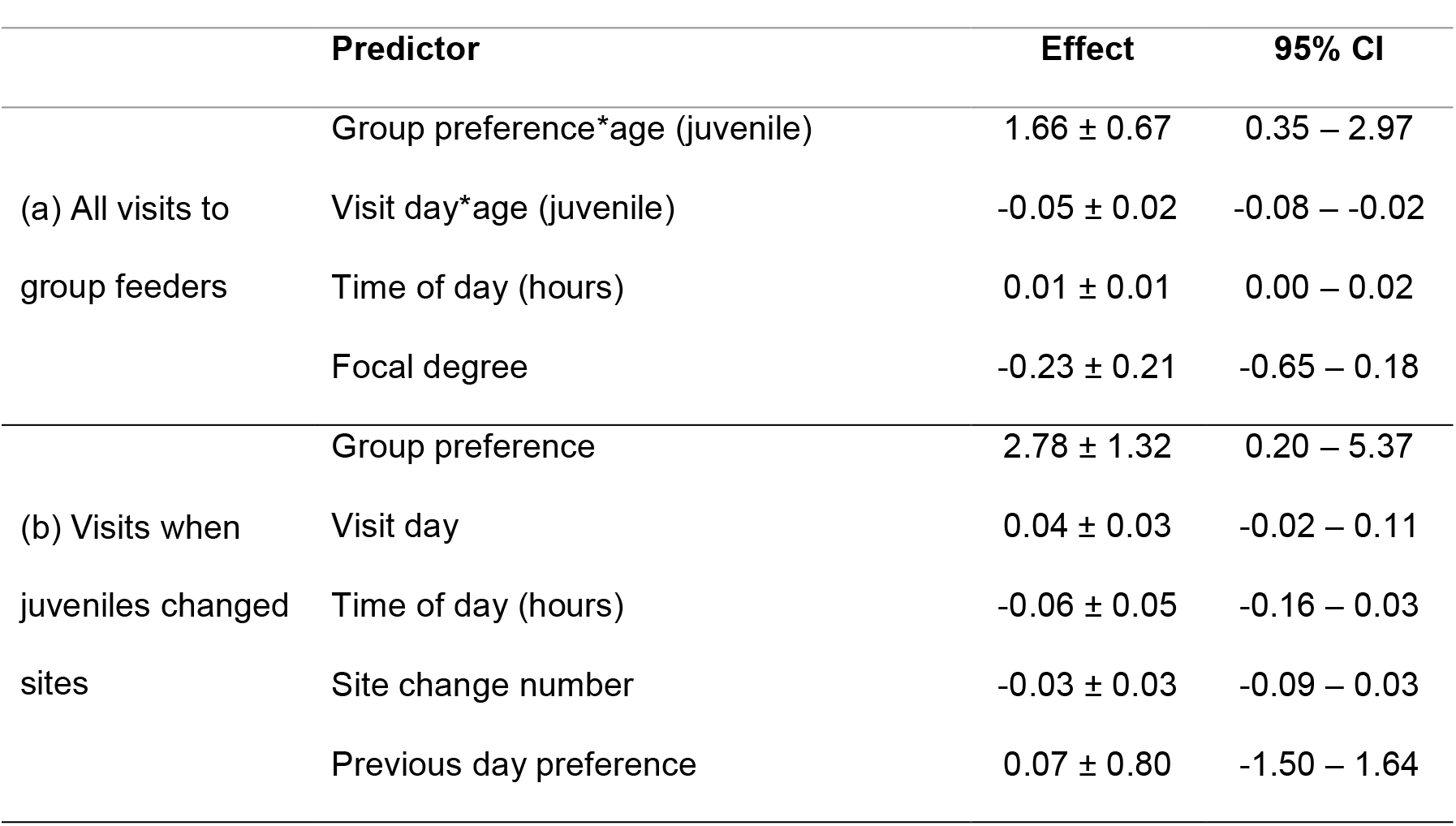
Effect sizes and 95% confidence intervals for predictors included in the top model set (ΔAICc < 2) analysing likelihood that hihi chose the locally-preferred side for (a) all visits and (b) when juveniles changed sites. Group preference = proportion end-preferred side was used by other hihi that day; focal degree of visiting bird calculated from the social network collected prior to side choice feeder setup.

After changing sites, juveniles also responded to the preferences of the new group. Juveniles were more likely to choose the local side when it was used most by the other group that day (Figure 5d, Table 3b, Supplementary Table 4b). This did not, however, vary with previous personal experience: side choice was not influenced by a stronger preference for the opposite side at their previous site (Table 3b, Supplementary Table 4b), and there was no effect of visit day or the number of times they had changed site (Table 3b). Time of day now had no effect on side choice (Table 3b; Supplementary Table 4b). Juveniles likely had the opportunity to observe multiple birds to assess group preference between leaving one site and using the feeder in the next, as the time between visiting feeders in different sites was longer than one hour in 70% of all site changes (median = 1.8 hours: IQR = 15.8; Supplementary Figure 1a) and a null GLMM model investigating variation in inter-site times was ranked higher than a model including experiment day (Supplementary Figure 1b; Supplementary Table 5a). Juveniles changed sites more times as the experiment progressed (effect of experiment day on number of changes per day = 0.06±0.01, 95% CI = 0.03 – 0.08; Supplementary Figure 1c; Supplementary Table 5b), so they could have encountered peers at both sites, multiple times per day. Finally, juveniles rarely followed an individual that had also changed sites (10/242 site changes), suggesting they were integrating into a group and not moving in flocks between feeders.

Although side choice was affected by the group’s preferences, hihi did not appear to copy particular individuals. Very few birds had a high (or low) probability of being copied, and most individuals were copied on approximately half their visits (Figure 5e; mean proportion copied = 0.50; 74% of individuals fell within 1 standard deviation of the mean). Adults and juveniles were equally likely to copy (included in models with ΔAICc < 2 but effect = –0.07 ± 0.10; 95% CI = –0.26 – 0.12) and did not change their behaviour if they were more social, according to the age of the previous individual, nor if they were more familiar with that preceding bird (Supplementary Table 6a). Instead, the only significant predictors were time between visits, as individuals became less likely to copy the previous bird when their visits were closer together (effect size of increasing time since preceding bird’s visit = 0.24 ± 0.04; 95% CI = 0.16 – 0.31; Supplementary Table 6a), and a stronger group preference that day (effect size = 0.65 ± 0.19; 95% CI = 0.27 – 1.03). The effect of time was stronger if the previous hihi was still at the feeder, as we re-analysed the top models but only included visits fewer than 30 seconds apart (Figure 5f; median time hihi spent inside feeders during 345 observations of feeding visits = 30s, IQR = 0; effect = 0.54±0.13, 95% CI = 0.28 – 0.80). Similarly, whenever juveniles changed sites no predictor significantly affected their likelihood of copying (null model ranked highest; Supplementary Table 6b).

## Discussion

Here, we demonstrated that social experiences with parents and peers in early life affected the foraging behaviour of young passerine birds. When given a choice between accessing a feeder on the left or on the right-hand side, recently-fledged hihi chose the same side as their parents. Furthermore, once they became independent and formed groups, juveniles that had previously encountered feeders with parents used a novel feeder before naïve birds. However, when encountering a feeder with a similar side-choice (but both sides were now equally rewarding), they did not maintain their parents’ preference. Instead, choice was initially random but over time all juveniles updated their choice in response to their peers’ behaviour. Juveniles paid more attention to this social information than adults did, but did not copy particular individuals. Consequently, the frequency of visits to one side of the feeder increased across the experiment, and to opposite sides at two different sites. Finally, individuals that switched between sites were more likely to match the locally-preferred option as more local birds chose that side.

Some behaviours are transmitted from parents to offspring directly through imitation or teaching (Thornton, 2006; van de Waal, Bshary and Whiten, 2014; Iwata *et al*., 2017) but time with parents can also facilitate learning new behaviour in other ways, such as promoting individual trial-and-error learning (Truskanov and Lotem, 2017). In our experiment, early-life encounters during parental care influenced how quickly juveniles used feeders once they became independent, but not how they interacted with the stimulus (their side choice at feeders was random). Furthermore, feeder juveniles’ responses were not determined by parents’ use of feeders at their nests. This suggests young hihi did not need to directly observe parent behaviour during early life, but instead they learned a generalised stimulus (feeder) response. Similar results have also been found in parrots (*Amazona amazonica*), where interactions with parents did not determine later responses to objects, but the presence of these objects during parental care did (Fox and Millam, 2004). Thus, lasting effects of time with parents may be subtle, and changes in behaviour can result from indirect influences.

By contrast, we found that young hihi copied their peers once independent while adults did not respond to the behaviour of others. Due to their inexperience, juveniles may rely more than adults on social information to determine behaviour (Laland, 2004; Kendal *et al*., 2005; Kendal, Coolen and Laland, 2009). Further, young hihi also altered their choices based on the overall collective behaviour of the group rather than specific individuals, as there was little evidence that familiarity and sociality affected hihi behaviour. Although individual-level familiarity or number of associates is important for information dissemination in some contexts (Aplin *et al*., 2012; Atton and Galef, 2014; Guillette, Scott and Healy, 2016; Ramakers *et al*., 2016), they may be less crucial when using collective information provided by groups. Generally, copying the predominant behaviour of groups rather than specific individuals is thought to help animals overcome some pitfalls of using social information (King and Cowlishaw, 2007; Conradt and Roper, 2005; but see Giraldeau, Valone and Templeton, 2002), such as the risk of copying misinformed individuals (Curio, Ernst and Vieth, 1978; Ward and Zahavi, 2008; Pruitt *et al*., 2016). If juveniles also have limited experience to judge reliability from different sources, then groups may provide a more complete picture of behavioural responses to the current environment (Clark and Mangel, 1984; Conradt and Roper, 2005). This could explain why we found little evidence that familiarity or degree, both specifically determined from visits to the feeders, affected hihi behaviour.

When paying attention to the collective behaviour of peers, animals may respond to small changes in the frequency of behaviour in groups to alter their own preferences and conform. Although there has been debate surrounding how to define conformity (for discussion, see Aplin *et al*., 2015a; van Leeuwen *et al*., 2015, 2016; Whiten and van de Waal, 2016), a general rule is that conforming animals tend to prefer a common behavioural option in a group, even if they have experience of alternative options (de Waal, 2013). Within the past 10 years behavioural conformity has been suggested to occur in taxa from invertebrates to humans, which highlights a widespread tendency for animals to copy the behaviour of others (birds: Aplin et al., 2015b; invertebrates: Fürtbauer & Fry, 2018; fish: Pike & Laland, 2010; primates, including humans: Haun, van Leeuwen, & Edelson, 2013; van de Waal et al., 2013; van de Waal, van Schaik, & Whiten, 2017; but see van Leeuwen et al., 2013). In support of this, juvenile hihi developed a preference for one feeder side which was driven more strongly by the behaviour of others than personal learning. Previous studies have further suggested animals quickly conform to traditions in new groups (van de Waal, Borgeaud and Whiten, 2013), and we also found evidence that copying was localised to the extent that juvenile hihi switched their behaviour when moving between groups with opposite preferences.

We acknowledge that with only two replicates of the feeder experiment in groups, it is possible that side choice patterns could have developed as a result of environmental effects (for example, surrounding vegetation). However, several lines of evidence suggest this was unlikely. Firstly, initial side choice was random at both feeders, suggesting no effect of an immediate bias. Secondly, the progressive changes in side choice are similar (albeit in the opposite direction) at both feeders and it is unlikely that both feeders would followed similar changes if there was an effect of some environmental aspect. Furthermore, side choice did not appear to be habit-driven (Pesendorfer *et al*., 2009) as first side choices did not predict overall individual preference. Instead, juvenile hihi paid attention to peers even with their own experience of the non-preferred side and when the cost of choosing that side was small. Overall, this suggests that juvenile hihi were copying predominant behaviours to conform to the group preference and adjusted their choices quickly irrespective of prior experience. This is only the second study (as far as we know) to demonstrate conformity patterns in wild birds (Aplin *et al*., 2015b).

While conforming can perpetuate behavioural biases in populations under certain conditions, emerging evidence shows that when conforming becomes a suboptimal strategy, animals reduce their tendency to copy (Aplin, Sheldon and McElreath, 2017). We also found that hihi were less likely to copy when using feeders at very similar times, and instead used the equally-rewarding alternate side. This could be a way to avoid queuing before feeding themselves, and maintain an optimal foraging intake (Milinski, 1982; Krebs and Inman, 1992). Even when animals have a propensity to disregard their own experience and copy the behaviours of others, they still pay attention to small trade-offs between competitive interactions and social learning strategies (Laland, 2004). Conformity in natural populations where there is social information use, competition, and/or resources of similar payoff (as in our experiment) may never result in a strong sigmoidal relationship between frequency of behaviour and likelihood of copying seen in previous studies (Aplin *et al*., 2015a, 2015b). Flexible use of conformist strategies under different levels of payoffs has only just begun to be considered (van Leeuwen *et al*., 2013; Aplin, Sheldon and McElreath, 2017), but here we suggest using social information to conform may particularly benefit naïve animals in less competitive environments.

Social information use is thought to play a key role in shaping ecological processes contributing to population stability, such as finding food, avoiding predation, and disease transmission (Blanchet, Clobert and Danchin, 2010). Sociality can be important to survival of juveniles in some species (for example feral horses, *Equus callabus* (Nuñez, Adelman and Rubenstein, 2015)). Therefore, clarifying the importance of juvenile groups for behaviour may help understand what influences survival during conservations translocations (Cox *et al*., 2014). Young hihi are often translocated to new sites to establish or supplement populations, so if peers are important *in situ* on Tiritiri Matangi Island, they may become more important at new sites or in environments where the risks of learning alone increase (Kendal *et al*., 2005; Webster and Laland, 2008; Rendell *et al*., 2010). However, because young hihi copied group behaviour (and did not copy according to familiarity), this could suggest that the specific identities of peers are not crucial to help them survive the critical post-release period. Ultimately, copying groups not individuals could allow juveniles to conform quickly to new environments, if paying attention to whichever individuals are present at the time is a way of attaining information about the current environment (Hatch and Lefebvre, 1997; Ramakers *et al*., 2016). Testing how groups change following translocations will further help understand the value of the group as a whole to juveniles of threatened species.

